# Synthetic-evolution reveals that phosphoregulation of the mitotic kinesin-5 Cin8 is constrained

**DOI:** 10.1101/312637

**Authors:** Alina Goldstein, Darya Goldman, Ervin Valk, Mart Loog, Liam J. Holt, Larisa Gheber

## Abstract

Cdk1 has been found to phosphorylate the majority of its substrates in disordered regions. These phosphorylation sites do not appear to require precise positioning for their function. The mitotic kinesin-5 Cin8 was shown to be phosphoregulated at three Cdk1 sites in disordered loops within its catalytic motor domain. Here, we examined the flexibility of Cin8 phosphoregulation by analyzing the phenotypes of synthetic Cdk1-sites that were systematically generated by single amino-acid substitutions, starting from a phosphodeficient variant of Cin8. Out of 29 synthetic Cdk1 sites that we created, eight were non-functional; 19 were neutral, similar to the phosphodeficient variant; and two gave rise to phosphorylation-dependent spindle phenotypes. Of these two, one site resulted in novel phosphoregulation, and only one site, immediately adjacent to a native Cdk1 site, produced phosphoregulation similar to wild-type. This study shows that, while the gain of a single phosphorylation site can confer regulation and modulate the dynamics of the spindle, to achieve optimal regulation of a mitotic kinesin-5 motor protein, phosphoregulation has to be site-specific and precise.

## Introduction

Mitotic cell division is an essential and highly regulated process by which genomic information in the form of duplicated chromosomes is faithfully transmitted from mother to daughter cells. This process is mediated by the mitotic spindle, a microtubule (MT)-based structure that undergoes a distinct set of dynamic morphological changes with precise temporal and spatial regulation. One of the major coordinators of the mitotic events is the conserved cyclin-dependent protein kinase 1 (Cdk1). The mitotic functions of Cdk1 are activated by association with the B-type cyclins (Clb(1-6) in yeast). The levels of the six cyclins oscillate (Morgan & Roberts, 2002), thus helping to provide stage-specific functions for Cdk1 (Nasmyth, 1996). Cdk1-dependent phosphorylation of mitotic substrates is also affected by the activities of protein phosphatases such as the PP2A^cdc55^ (Queralt, Lehane et al., 2006) and Cdc14 (D’Amours & Amon, 2004, Stegmeier & Amon, 2004), which dephosphorylate Cdk1 target sites. Thus, mitotic events are coordinated by the balance between the activities of Cdk1 and opposing phosphatases.

Previous work identified a large class of “flexible” Cdk1 substrates that were enriched for phosphorylation sites throughout evolution, but where the precise site position was not strictly conserved (Holt, Tuch et al., 2009). These phosphorylation sites tended to occur in disordered regions, including loops and termini of proteins. It was suggested that these phosphates might interact with modular phospho-binding motifs or disrupt a protein-protein interface, forms of regulation that are relatively tolerant of changes in site position (Holt et al., 2009). A well-characterized example of a Cdk1 target with a flexible phosphorylation cluster is the Sic1 protein, a Cdk1 inhibitor in *Saccharomyces cerevisiae* (Mendenhall, Jones et al., 1987, Schwob, Bohm et al., 1994). For proper phosphoregulation of Sic1, a minimal set of priming phosphorylation sites and phospho-degrons is required, while distances between the sites in the cluster can be flexible (Koivomagi, Ord et al., 2013, Koivomagi, Valk et al., 2011a). A second, smaller set of substrates was found that maintained a precise position throughout long periods of evolution. These sites were often found in ancient metabolic enzymes, suggesting that they may have evolved early and adopted a highly efficient form of phosphoregulation that relies on exact conformation changes (Holt et al., 2009). To date, there is very little experimental data directly testing the flexibility of phosphoregulation.

To address this issue, we experimentally examined the flexibility of phosphoregulation by Cdk1 using the *S. cerevisiae* mitotic kinesin-5 Cin8 as a model-protein. These conserved bipolar kinesin-5 motors perform essential roles in mitotic spindle dynamics of eukaryotic cells by crosslinking and sliding apart antiparallel microtubules of the spindle (reviewed in (Ferenz, Gable et al., 2010, Goulet & Moores, 2013, Hildebrandt & Hoyt, 2000, Kashina, Rogers et al., 1997, Singh, Pandey et al., 2018, Valentine, Fordyce et al., 2006, Valentine & Gilbert, 2007, Waitzman & Rice, 2014)). Their function had been shown to be regulated by Cdk1 in various organisms (Avunie-Masala, Movshovich et al., 2011, Blangy, Arnaud et al., 1997, Blangy, Lane et al., 1995, Cahu, Olichon et al., 2008, Chee & Haase, 2010, Crasta, Huang P Fau - Morgan et al., 2006, Gardner, Bouck et al., 2008, Goldstein, Siegler et al., 2017, Shapira & Gheber, 2016, Sharp, McDonald et al., 1999). The budding yeast *S. cerevisiae* encodes two kinesin-5 homologues, Cin8 and Kip1, that partially overlap in spindle assembly and maintenance of the bipolar spindle structure (Hoyt, He et al., 1992, Roof, Meluh et al., 1992, Saunders & Hoyt, 1992), in focusing kinetochore clusters (Gardner et al., 2008, Tytell & Sorger, 2006, Wargacki, Tay et al., 2010), in anaphase B spindle elongation (Saunders, Koshland et al., 1995, Straight, Sedat et al., 1998), and in stabilizing and organizing the middle-spindle midzone, an overlapping array of antiparallel MTs (Fridman, Gerson-Gurwitz et al., 2009, Fridman, Gerson-Gurwitz et al., 2013, Gerson-Gurwitz, Movshovich et al., 2009, Ibarlucea-Benitez, Ferro et al., 2018, Movshovich, Fridman et al., 2008).

Kinesin motors undergo diverse phosphoregulation, including by Cdk1 (reviewed in (Gibbs, Greensmith et al., 2015, Goulet & Moores, 2013, Waitzman & Rice, 2014)). In particular, the mitotic functions of both *S. cerevisiae* kinesin-5 Cin8 and Kip1 were shown to be regulated by Cdk1 (Avunie-Masala et al., 2011, Chee & Haase, 2010, Goldstein et al., 2017, Shapira & Gheber, 2016). For Cin8, such phosphoregulation occurs primarily at three Cdk1 sites, which are located in disordered loops 8 and 14 within the catalytic motor domain (Avunie-Masala et al., 2011, Goldstein et al., 2017, Shapira, Goldstein et al., 2016). We therefore asked how easy it is to recapitulate or create new phosphoregulation by generating novel Cdk1 sites within Cin8, adjacent to and distant from the native sites. To answer this question, we examined the phenotypes of 29 new possible Cdk1-sites that were systematically generated by a single amino-acid substitution starting from a phosphodeficient allele of Cin8. By combining a comprehensive genetic, cell biological and biochemical characterization of these mutants we found that the three native sites in the catalytic domain of Cin8 can each alone support partial regulation of Cin8. We also found that out of 29 novel synthetic Cdk1 sites that we created, eight (28%) resulted in non-functional Cin8, 19 (65%) resulted in a neutral spindle phenotype similar to the phosphodeficient variant, and two (~7%) gave rise to phosphorylation-dependent spindle localization phenotypes. Of these two, one site, one amino-acid proximal to a native Cdk1 site, closely recapitulated the original phosphoregulation without destabilizing Cin8 or perturbing its function. This study shows that while the gain of a single phosphorylation site can confer complex regulation and thereby modulate the dynamics of the mitotic spindle, to achieve optimal regulation of a mitotic kinesin-5 motor protein, phosphoregulation has to be highly constrained.

## Results

### Strategy for generation of synthetic Cdk1 phosphorylation sites in Cin8

Our strategy to examine the flexibility of phosphoregulation of Cin8 by Cdk1 was to introduce novel consensus Cdk1 sites in the Cin8 ORF in a systematic manner. As a basis for mutagenesis, we used a phosphodeficient variant of Cin8 (Cin8-5A) carrying mutations of the phosphoacceptor serines and a threonine to alanine within the five native Cdk1 sites of Cin8, at positions S277, T285, S493, S763, and S1010. New Cdk1 sites were systematically created by single amino-acid replacement in most locations such that replacement resulted in a novel partial or full Cdk1 consensus site (Fig. 1A and B). We comprehensively created mutations around every proline as this amino acid is the major specificity determinant for Cdk1. For every proline, a serine was introduced at position -1 to generate a Cdk1 phosphorylation consensus sequence (S-P or S-P-x-K/R). If lysine or arginine were already present at the +3 position relative to the new serine, this mutation immediately generated a full site. Otherwise, a partial site was created (also referred to as a minimal consensus site). In these cases, a second synthetic allele was additionally generated with a lysine at position +3 to the serine, to produce the full Cdk1 phosphorylation consensus [S-P-x-K] (Fig. 1B class 1). In addition, we created Cdk1 sites by substitution of a proline only in cases where full Cdk1 phosphorylation consensus sites were created by this single change (i.e., [S-x-x-K] to [S-P-x-K]; Fig. 1B class 2). We limited the proline substitutions in this fashion because proline substitutions are often poorly tolerated due to the structural constraints of this amino acid. To avoid any bias, we generated the new sites based on the sequence only, without excluding structured and conserved regions. Using this approach, we both reintroduced three native Cdk1 sites at the motor domain of Cin8, termed Cin8 natural-phosphorylation variants (Cin8_nat_), and additionally generated 29 novel sites termed Cin8 synthetic-phosphorylation variants (Cin8_syn_). For variants that exhibited potential phosphoregulation phenotypes (see below), we created control mutations in which the serine within the consensus sequence was changed to alanine, which lacks a phosphoacceptor hydroxyl group (Fig. 1B) (see Materials and Methods).

**Figure 1:**
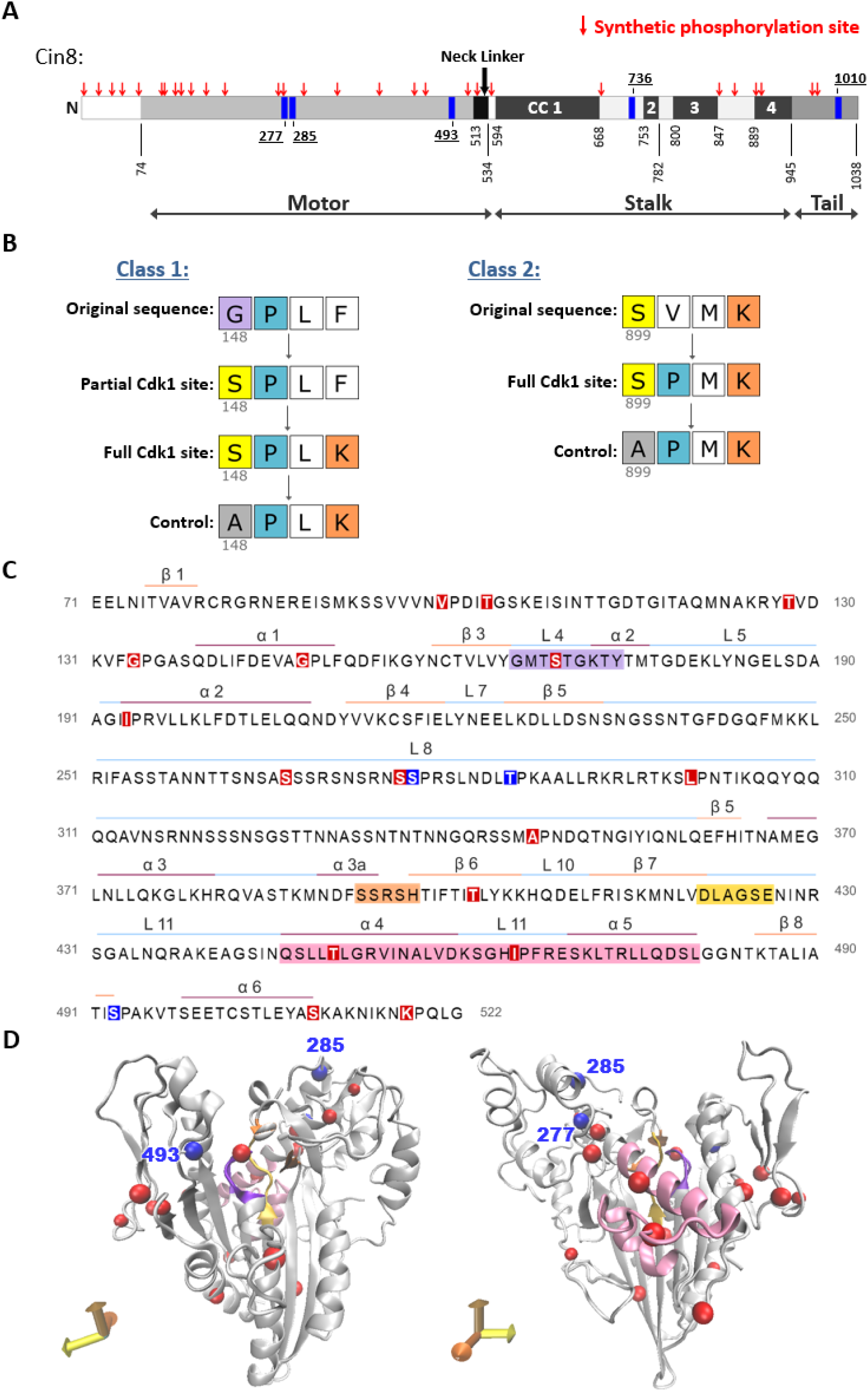
A strategy to generate synthetic Cdk1 sites. (A) Schematic representation of Cin8. Native Cdk1 phosphorylation sites are indicated by blue rectangles, synthetic phosphorylation sites are marked by red arrows. CC: coiled coil. (B) Mutagenesis strategy to create the synthetic phosphorylation sites. Class 1 synthetic sites are more structurally conservative; an amino acid prior to an existing proline was mutated to serine to create a minimal Cdk1 consensus site. For synthetic sites that didn’t result in full Cdk1 phosphorylation consensus by a single serine insertion, we also generated a full consensus site by additional mutation of the position +3 relative to the potential phosphoacceptor serine to lysine. Finally, control mutants were generated for synthetic sites, which resulted in Cin8 detachment from the spindle during spindle elongation by mutating the serine at the phosphoacceptor position to alanine; see Materials and Methods. Class 2 synthetic sites are more structurally perturbative because the amino-acid proline was introduced at +1 relative to an existing serine or threonine with a lysine or arginine at the +3 position, resulting in a full consensus Cdk1 site. As in class one, an alanine control mutant was generated in some cases. (C) Sequence of Cin8 motor domain with secondary structure annotated according to (Subbiah & Harrison, 1989). Key domains: ATPase p-loop in purple, Switch I in orange, Switch II in yellow, MT biding domain (helix 4, loop 12 and helix 5) in pink. Native and Synthetic Cdk1 phosphorylation sites are indicated by blue and red, respectively. (D) Model of Cin8 motor domain predicted by Swiss-model based on human kinesin-5 Eg5 (PDB ID: 3HQD) presented in two different views indicated by axis arrows on the bottom left. Cdk1 sites represented by spheres, color coded as in C, blue for native and red for synthetic sites.

### Single Cdk1 sites at the native positions confer regulation

Re-generation of the native Cdk1 sites by the above described approach allowed us to examine the functionality of these sites as a sole source of Cdk1 phosphoregulation and compare their phenotypes to the wt Cin8 and the phosphodeficient Cin8-5A. We previously demonstrated that each of the native Cdk1 phosphorylation sites in the motor domain of Cin8, S277, T285, and S493, are independently phosphorylated by Cdk1 *in vitro* and that mutations to alanine of each of these sites cause different phenotypes during anaphase (Goldstein et al., 2017). Here, the native sites were examined as partial (suffixed by “p”) and full (suffixed by “f”) Cdk1 consensus sequences since they were generated as variants of class 1 (Fig. 1B). It is worth noting that although the native Cdk1 phosphorylation site at position 285 is composed of a threonine followed by a proline, here we introduced a serine to result in S-P consensus (see Materials and Methods). The native variants were first examined by general viability test, then by two dynamic spindle-localization features: translocation to the midzone at mid-anaphase and detachment from the spindle at late anaphase. Finally, the functionality of the variants was evaluated by analysis of anaphase spindle elongation dynamics (Goldstein et al., 2017).

In order to determine if modifications to the native sites substantially alter the functionality of Cin8, we conducted a yeast viability test in which the Cin8_nat_ variants were expressed from a centromeric (CEN) plasmid as a sole source of kinesin-5 function in cells deleted for the chromosomal copies of *CIN8* and *KIP1*. Cin8 and Kip1 have overlapping roles. However, at least one of them is required for proper spindle formation and yeast viability (Avunie-Masala et al., 2011, Duselder, Fridman et al., 2015, Gheber, Kuo et al., 1999, Hoyt et al., 1992, Roof et al., 1992, Saunders & Hoyt, 1992). We found that cells expressing native partial and full Cin8 phosphorylation variants (Cin8_nat_) and their controls, in which the serine at the Cdk1 site was mutated to alanine, are viable at all temperatures (Fig. 2A). Therefore, these single phosphorylation site alleles are sufficient for the essential function of Cin8.

**Figure 2:**
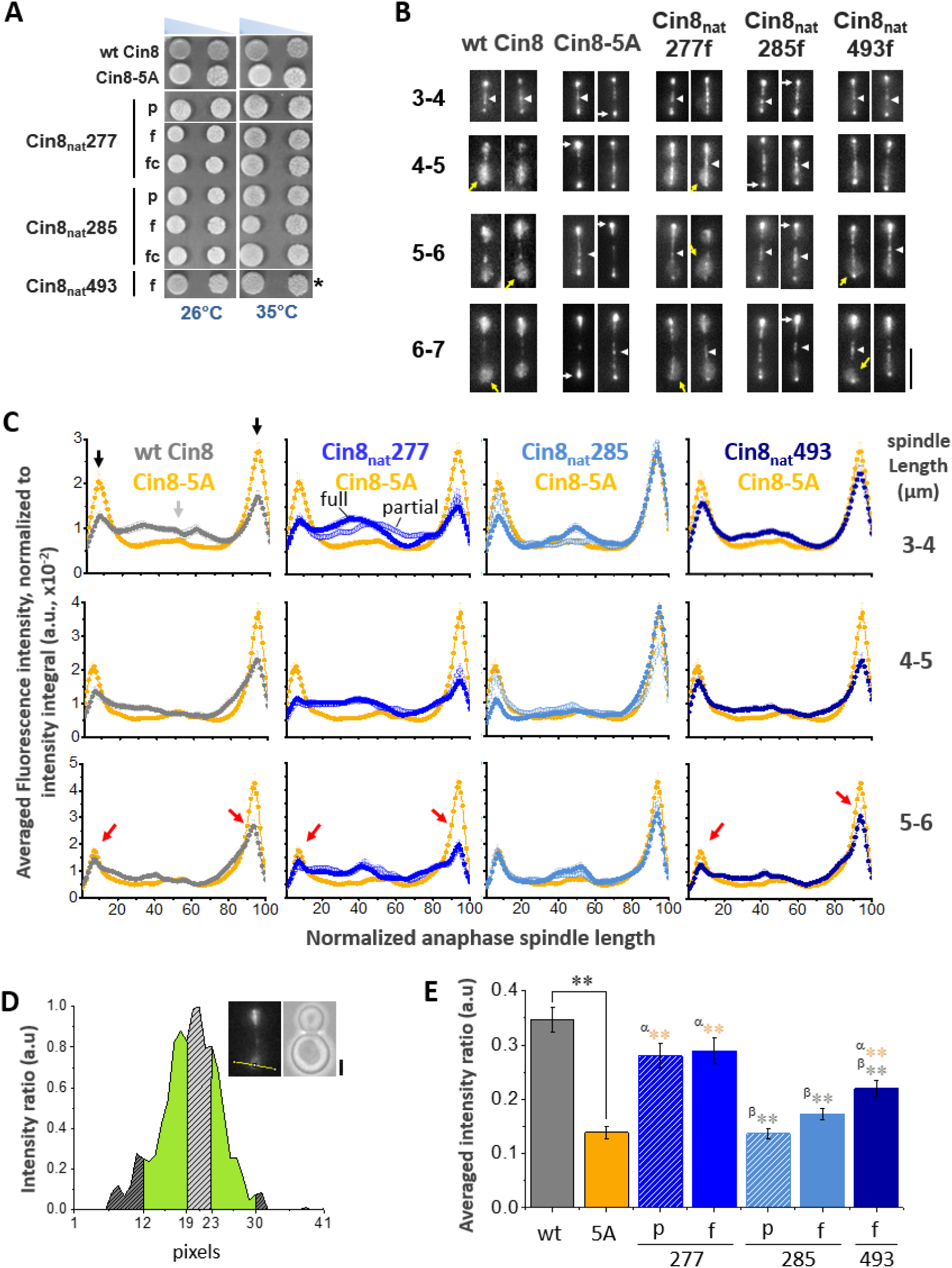
A single native phosphorylation site is sufficient for viability and each site regulates the distribution of kinesin-5 differently. (A) Viability test of a shuffle yeast strain deleted for its chromosomal copies of *CIN8* and *KIP1* and containing a recessive chromosomal cycloheximide resistance gene (*cin8Δkip1Δcyh^r^*) on plates containing 7.5 μg/ml cycloheximide. Cells contain centromeric (CEN) plasmids expressing the Cin8 phospho-variants with one native Cdk1 site as a sole source of Cdk1 phospho-regulation (indicated on the left, e.g., Cin8_nat_277 is a cin8Δ; kip1Δ strain with a plasmid containing Cin8 with only the CDK1 consensus site at position 277 intact). Cells were plated in serial dilution, grown at 26°C and 35°C indicated at the bottom; *grown at 37°C (Avunie-Masala et al., 2011, Duselder et al., 2015). (B) Localization of Cin8 native phospho-variants (Cin8_nat_) to anaphase spindles. Anaphase spindles are divided into length categories, indicated on the left. Representative 2D projection of a 3D stack of fluorescence images of cells expressing 3GFP-tagged Cin8 phospho-variants from a CEN plasmid, indicated on the top. All cells are presented with mother cell at the bottom. Scale bar is 5 μm. Yellow arrows indicate Cin8 detachment from the SPBs, white arrows indicate Cin8 accumulation at the SPBs and white arrowheads indicate Cin8 distribution at the midzone. (C) Fluorescence intensity along the spindle of the wt Cin8 and Cin8_nat_ mutants, compared to the fluorescence of Cin8-5A (orange). Cin8-3GFP fluorescence intensity along to the spindle was measured starting at the mother pole and divided to 100 segments of equal length. The signal was normalized to the integral of spindle-bound Cin8-3GFP signal. The signal for each mutant was compared to that of Cin8-5A, indicated on top. Spindles were binned into length-ranges indicated on the right. Full and partial Cdk1 consensus sequences are indicated by filled and hollow circles, respectively. Black arrows indicate Cin8 accumulation at the SPBs. Light gray arrow indicates elevated distribution of Cin8 at the midzone region. Red arrows indicate reduced Cin8 distribution at the SPBs compared to Cin8-5A. (D) Analysis method to quantify Cin8 detachment from the spindle pole. A 40-pixel line was created perpendicular to the spindle at the mother SPB. The *Y*-axis represents the intensity measured along the yellow line, following subtraction of the background and normalized to the maximum intensity. The corresponding image of a representative cell is shown at the top right; scale bar: 2 μm. The brightest 6 pixels were assigned as Cin8 attached to the center of the SPB and are indicated on the graph as dashed gray area. Intensity values of all pixels are normalized to the maximal intensity, usually at the SPB. The 11 distal pixels on the far ends of the line (which are outside the anaphase nucleus) were averaged and subtracted as background (dark gray dashed area). The fluorescence intensity of the 6 pixels immediately flanking the SPB was assigned Cin8 detached from the spindle (green area). (E) Average intensity ratio of the Cin8 detachment at spindle length of 6-7 μm, as explained in D, of Cin8_nat_ mutants compared to wt Cin8 and Cin8-5A. Average ± SEM of 18-20 cells is shown. Significance was determined by post-ANOVA all-pairwise comparison using Tukey procedure. α (orange): *p*-value compared to Cin8-5A, β (grey): *p*-value compared to wt Cin8. \**p* < 0.05, \*\**p* < 0.01. (A)-(E): p - partial phosphorylation consensus (S-P), f - full phosphorylation consensus (S-P-x-K), fc - full control phosphorylation consensus (A-P-x-K).

Next, we examined the localization of Cin8 variants, tagged with 3GFP, during anaphase B spindle elongation (Fig. 2B). Consistent with previous results (Avunie-Masala et al., 2011, Goldstein et al., 2017), we observed that at the early stages of anaphase Cin8 is distributed along the spindle and later detaches from the spindle and becomes diffusively localized around the spindle pole bodies (SPBs), likely in the divided nuclei (Fig. 2B). In order to quantitatively compare the localization patterns of Cin8 variants during anaphase B, we performed line-scan intensity analysis of Cin8 distribution along the spindle at different anaphase spindle lengths, as previously described (Goldstein et al., 2017). We found that the phosphodeficient mutant Cin8-5A is mainly concentrated at the SPBs (Fig. 2C, black arrows), while wt Cin8 exhibited a lower concentration at the SPBs and higher levels along the spindle (Fig. 2C gray arrow). This result indicates that Cin8-5A is unable to translocate from the SPBs to the spindle midzone and that translocation is regulated via phosphorylation of Cin8 in at least one Cdk1 site. We previously obtained similar results with the Cin8-3A variant, carrying phosphodeficient mutations to alanine at the three Cdk1 sites in the catalytic domain of Cin8 (Goldstein et al., 2017). This indicates that, consistent with a previous report (Avunie-Masala et al., 2011), phosphoregulation of Cin8 is dependent mainly on phosphorylation of Cdk1 sites in its catalytic domain. Quantitative analysis of spindle localization also revealed that in short spindles of 3-4 μm, the partial and full Cin8_nat_277 variants and the full Cin8_nat_493f but not the partial Cin8_nat_285p had recruitment of Cin8 throughout the spindle length (Fig. 2C 3- 4 μm), similarly to wt Cin8. In contrast, the full Cin8_nat_285f exhibited a sharp peak of Cin8 at the middle of the spindle (Fig. 2C 3-4 μm). Overall, our data indicate that the S277 and S493 sites present phenotypes that are similar to the wt Cin8, while S285 site is similar to the Cin8-5A variant, consistent with our previous report (Goldstein et al., 2017).

To characterize the detachment of the native Cin8 phosphorylation variants from the spindle at late anaphase (Fig. 2B, yellow arrows), we performed quantitative analysis of Cin8 fluorescence signal perpendicular to the spindle (Fig. 2D) as previously described (Goldstein et al., 2017). We found that the detachment of wt Cin8 from the spindle in late stages of anaphase B was abolished by the phosphodeficient mutant Cin8-5A (Figs. 2E and EV3), indicating that phosphorylation by Cdk1 is required for this detachment (Avunie-Masala et al., 2011, Goldstein et al., 2017). We found that partial and full native variants at position S277 exhibit high degree of detachment, most similar to that of the wt Cin8; partial and full variants at position S285 being no different from the phosphodeficient CIn8-5A, while the variant at position S493 has an intermediate effect (Figs. 2E and EV3). This is consistent with previous reports that phosphorylation at the S277 site is the major regulator of Cin8 distribution along the spindle and its detachment from the spindle.

### Categorization of the synthetic Cdk1 sites

Results presented so far demonstrate that single Cdk1 sites in the motor domain of Cin8 produce phenotypes markedly different from those of the phosphodeficient Cin8-5A (Fig. 2), indicating that phosphorylation at a single Cdk1 site can, at least partially, regulate the function of Cin8. Thus, we next examined the phenotypes of the synthetic phospho-variants of Cin8 (Cin8_syn_). Since phosphoregulation among kinesin-5 motors produces varied effects including effects on motor activity (Gibbs et al., 2015, Goldstein et al., 2017, Morfini, Schmidt et al., 2016, Ritter, Kreis et al., 2015, Waitzman & Rice, 2014, Wojcik, Buckley et al., 2013), predicting possible outcomes as a result of introducing novel sites for Cdk1 phosphorylation is not trivial. Yet we postulated hypothetical outcomes: we expect some mutations to result in loss of function due to perturbation of Cin8 structure. Other mutants may result in no structural perturbation and no observable phosphoregulation effect; these would be neutral mutations. And finally, there may be mutants that result in regulation of Cin8 function, either by recapitulating the original phosphoregulation or resulting in novel regulation. To distinguish between these possibilities, the 3GFP-tagged Cin8_syn_ variants were first screened for their localization to anaphase spindles by live-cell imaging (Fig. 3) and for their ability to support the viability of cells deleted for the function of Cin8 and Kip1 as a sole source for kinesin-5 function (Fig. 3). Out of the 29 Cin8_syn_ mutants that were generated, eight (28%) were found to carry deleterious mutations and resulted in cell death. These mutants exhibited mostly diffusive detachment of Cin8 from the spindle that was not abolished by the alanine control variants and thus were not due to phosphorylation (Fig. 3). Of these eight variants, five carried proline insertion mutations (Fig. 1B Class 2), likely disrupting the secondary structure of Cin8. The largest class of mutants were neutral. We found 19 mutants (~65%) that had no discernible effect on localization, i.e., Cin8-5A-like phenotype (Fig. 3) (see Materials and Methods). These sites could be neutral for several reasons. They may not be accessible for Cdk1 phosphorylation, phosphatases may keep steady-state phosphorylation levels low, or it could be that phosphorylation at these sites does not influence Cin8 function. Finally, two novel Cdk1 positions (7%) resulted in a phosphorylation-dependent spindle-localization phenotype, i.e., control variants abolished or corrected part of the phenotypes induced by the novel Cdk1 sites S148 and S276 (Figs. 3, 4, EV1, EV2, and EV3). The last were further investigated by quantitative dynamic localization analysis (Figs. 4 and EV3), *in vitro* kinase assays, and spindle elongation measurements (Figs. 5, 6, and EV6).

**Figure 3:**
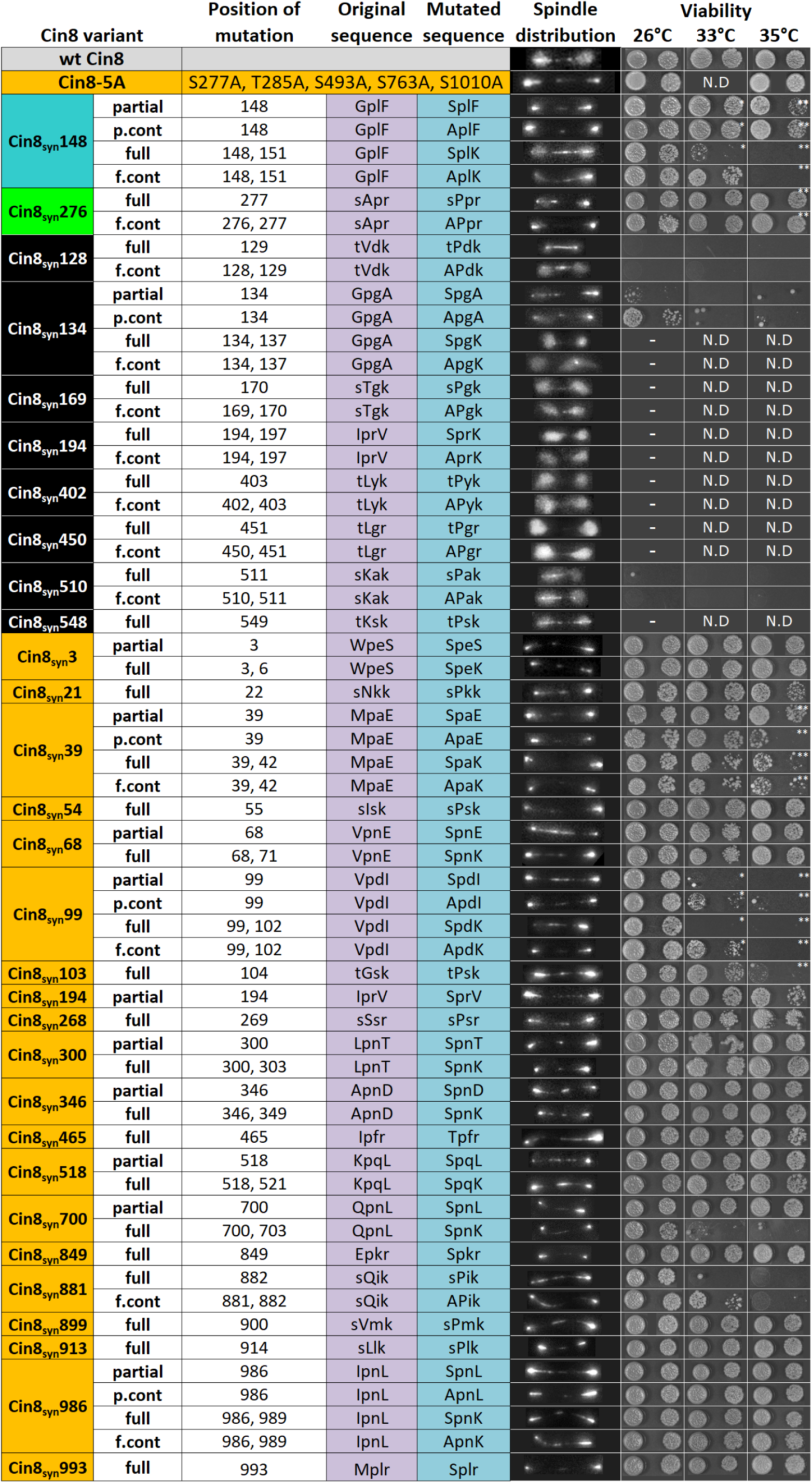
Phenotypes of all the Cin8_syn_ mutants examined in this study. The name and the Cdk1 phosphorylation consensus are indicated on the left (partial [S/T-P], partial control (p.cont) [A-P], full [S/T-P-x-K/R], and full control (f.cont) [A-P-x-K/R]), color coded: wt Cin8 (gray), Cin8-5A, and mutants exhibiting no detachments from the spindle (orange), mutants that exhibited reduced viability, or are non-viable at 26°C (black), Cin8_syn_148 (cyan), and Cin8_syn_276 (green). In the middle: position of mutations, original sequence (purple) and mutated sequence (blue) capital letters represent positions of mutations. Distribution of Cin8-3GFP along the spindles and its detachment (spindle distribution) are shown following the mutated sequence. Viability test results at 26°C, 33°C, and 35°C are shown on the right. * the temperature was 34°C, ** the temperature was 37°C. N.D. – not determined.

**Figure 4:**
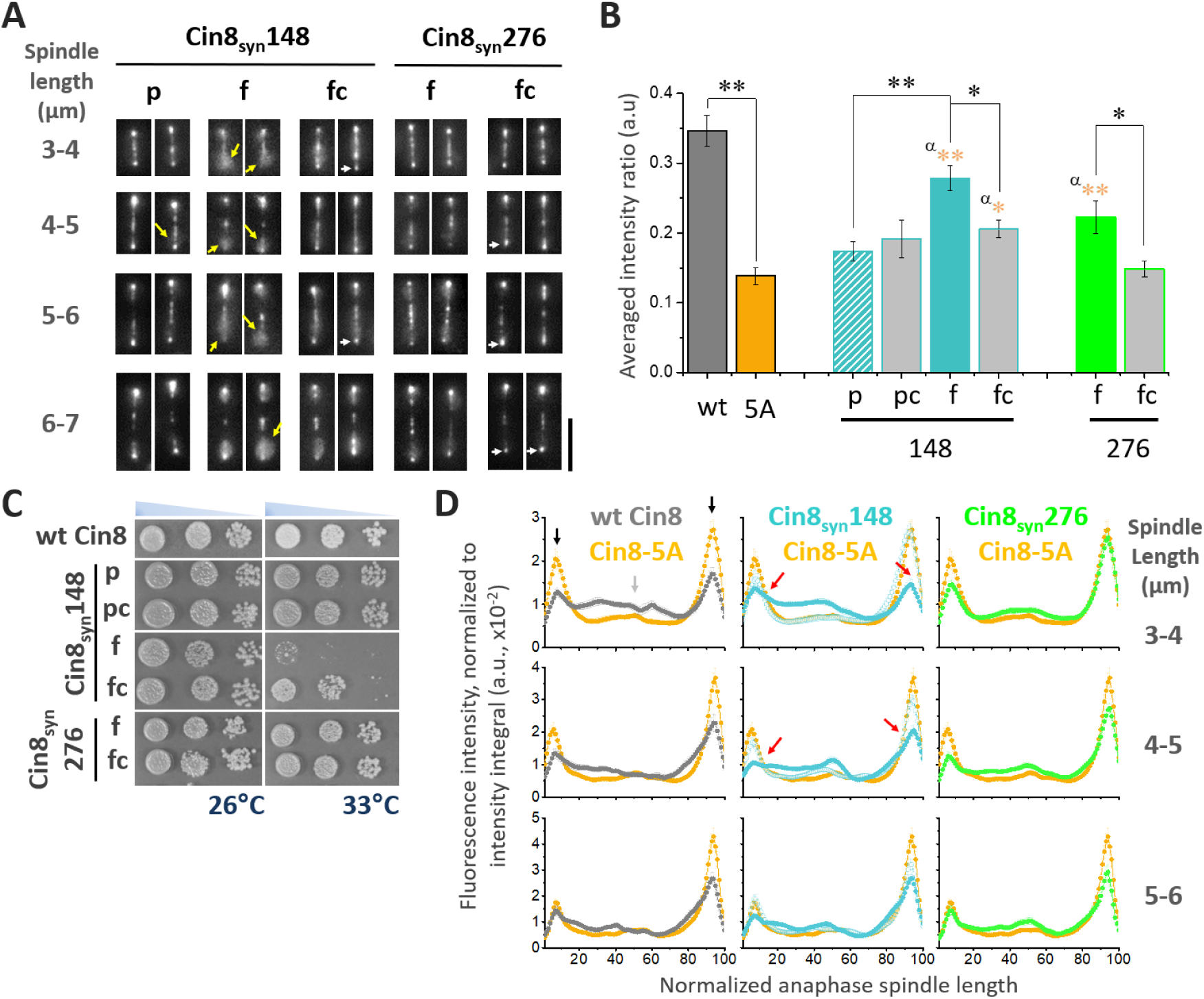
*In vivo* phenotype of selected synthetic Cin8 variants. (A) Representative 2D projection of a 3D stack of fluorescence images of cells expressing 3GFP-tagged Cin8_syn_ phospho-variants, indicated on the top. Spindle lengths are indicated on the left. All cells are presented with the mother cell at the bottom. Yellow arrows indicate Cin8 detachment from the spindle; white arrows indicate Cin8 concentration at the SPBs. Scale bar: 5 μm. (B) Intensity ratio perpendicular to the spindle (as in Fig. 2D and E) of Cin8_syn_ variants, compared to wt Cin8 and Cin8-5A. Average ± SEM of 18-20 cells (except Cin8_syn_148pc – see Materials and Methods) at spindle length of 6-7 μm is shown. Significance was determined post-ANOVA all-pairwise comparison using Tukey procedure. α: *p*-value compared to Cin8-5A. All variants except Cin8_syn_148f exhibited reduced detachment from the spindle compared to wt Cin8 (*p* < 0.01). p: partial Cdk1 consensus sequence [S/T-P], pc: control of partial consensus sequence [A-P], f: full Cdk1 consensus sequence [S/T-P-x-K/R], fc: control of full consensus sequence [A-P-x-K/R]. \**p* < 0.05, \*\**p* < 0.01. (C) Viability of a shuffle *cin8Δkip1Δcyh^r^* strains as in Fig 2A expressing Cin8_syn_ mutants. Cells were plated in serial dilution and grown on YPD plates with 7.5 μg/ml cyclohexamide at 26°C and 33°C. (D) Average fluorescence intensity along the spindle of Cin8_syn_ mutants, indicated on top, compared to the fluorescence of Cin8-5A as in Fig. 2C. Black arrows point to Cin8 accumulation at the SPBs. Light gray arrow indicates elevated distribution of Cin8 at the midzone region. Red and green arrows indicate reduced distribution at the SPBs and accumulation at the midzone compared to Cin8-5A, respectively.

**Figure 5:**
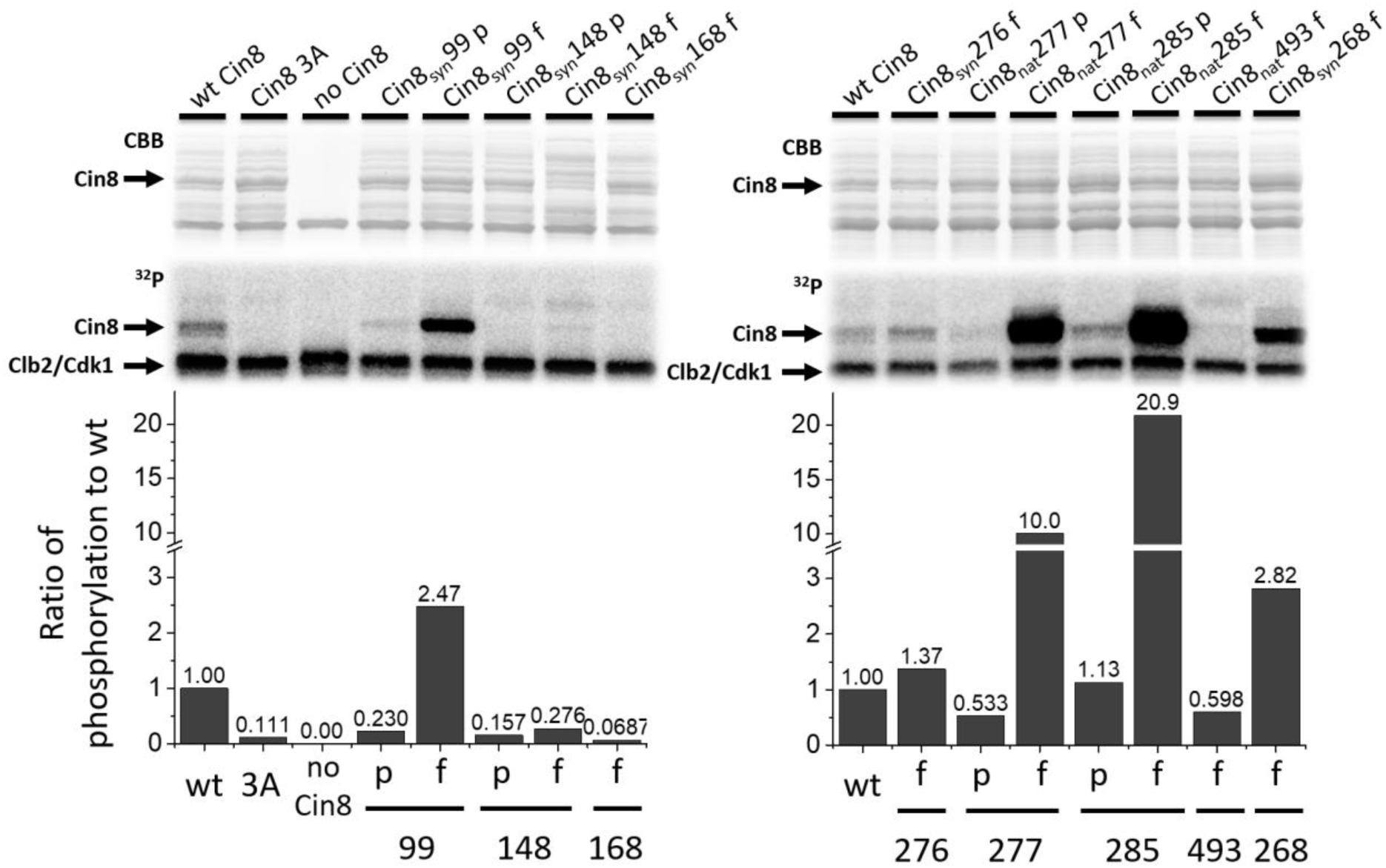
Synthetic sites are phosphorylated by Cdk1 *in vitro*. *In vitro* Cdk1/Clb2 kinase assay of the Cin8 motor domain. Top – Coomassie brilliant blue staining (CBB) and ^32^P autoradiograms of SDS-PAGE fractionation of phosphorylation reaction mixtures with bacterially expressed motor domain of wt Cin8, phosphodeficient variant Cin8-3A; and phospho-variants of Cin8. The phosphorylation reaction mixture without Cin8 served as negative control (no Cin8). Bottom: qualitative level of phosphorylation. The columns quantify phosphorylation band intensity, as measured on the autoradiogram and normalized to the amount of Cin8 as derived from CBB gel. The columns are named after the position of the phosphorylation and the variation of the Cdk1 phosphorylation consensus; “p” for partial and “f” for full.

**Figure 6:**
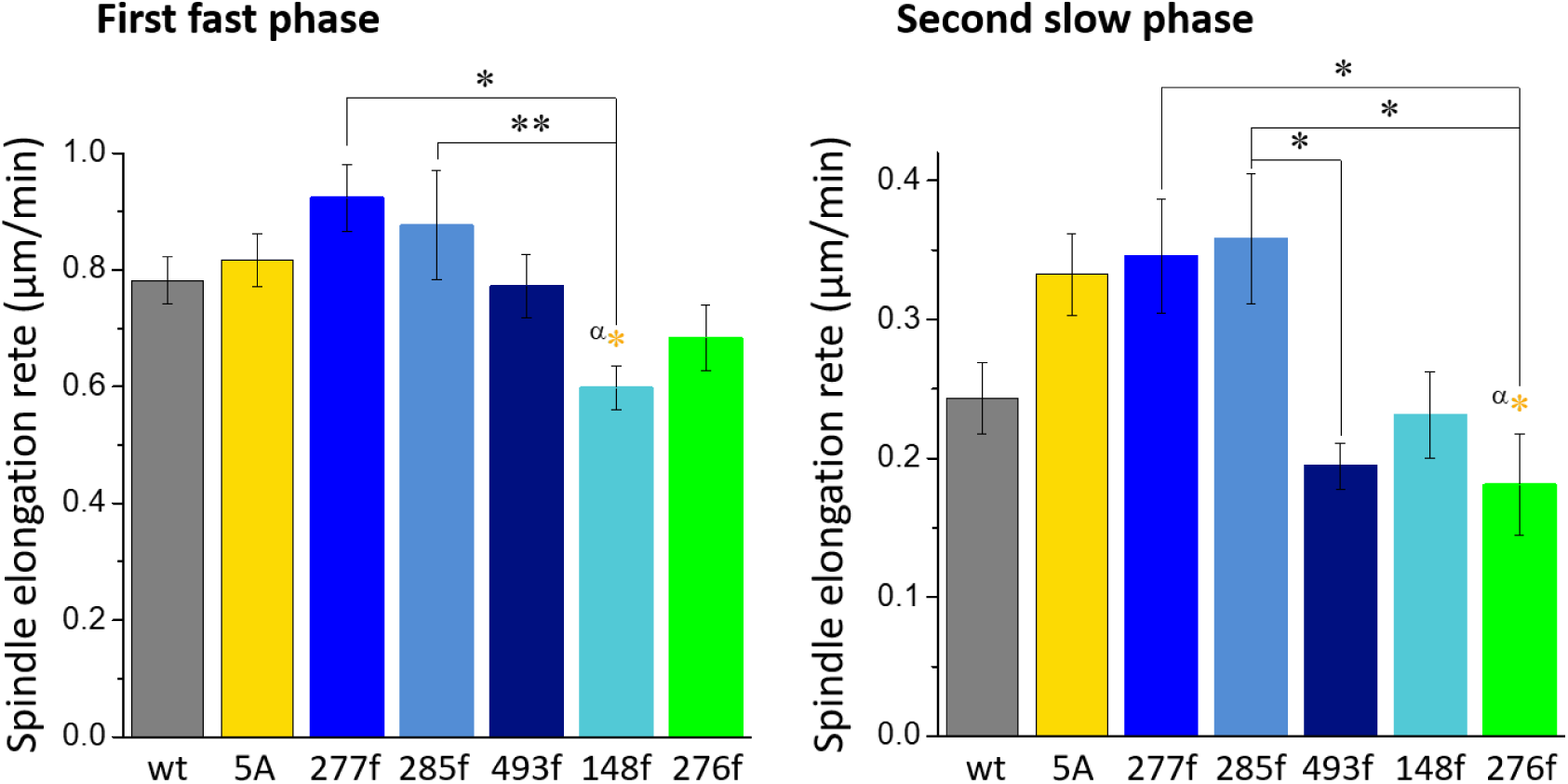
Synthetic and native phospho-variants differently affect anaphase spindle elongation *in vivo*. Spindle elongation rates of the first-fast and the second-slow phases of the full native and synthetic variants. Averages ± SEM of 9-12 cells are shown. Phospho-variants of Cin8 are indicated on the bottom, 5A represents the phosphodeficient Cin8-5A. Significance was determined by post-ANOVA all-pairwise comparison using Tukey procedure. α (orange): p-value compared to Cin8-5A. \**p* < 0.05, \*\**p* < 0.01.

### Two synthetic Cdk1 sites confer differential phosphoregulation

The novel sites at positions S148 and S276 exhibited differential effects on the dynamic detachment of Cin8 from the spindle (Figs. 3, 4A, B, EV1, EV2 and EV3). The partial Cin8_syn_148p variant showed no detachment, similar to the control variant Cin8_syn_148pc. However, the full Cin8_syn_148f variant exhibited high levels of detachment from the spindle at anaphase, comparable to wt Cin8. This novel regulation was lost in the non-phosphorylatable alanine control allele Cin8_syn_148fc (Figs. 4A, B, EV1, and EV2), indicating that Cin8 detachment from the spindle of the full Cin8_syn_148f is dependent on phosphorylation at this site. The full Cin8_syn_276f variant also exhibited detachment from the spindle but to a lower extent. This detachment was also abolished by the control Cin8_syn_276fc mutation (Figs. 4A, B, EV1, and EV2), indicating that this detachment is also phosphorylation dependent.

Cin8_syn_276f, which contains a full Cdk1 phosphorylation consensus sequence in position S276 and is of Class 2 (Fig. 1B), was viable at all temperatures, similar to the wt Cin8. In contrast, the mutant bearing a full Cdk1 consensus sequence at position S148 (Cin8_syn_148f) was temperature sensitive at temperatures higher than 33°C (Figs. 3 and 4C). The control variant Cin8_syn_148fc partially rescued the phenotype and grew at 33°C (Fig. 3 and 4C), indicating that gain of phosphate at these positions interferes with Cin8 function. All S148 variants were stable at 26^°^C and 33^°^C, similarly to the native and the synthetic S276 variants (Fig. EV4), suggesting that mutation at this site affects mainly protein function and not its stability.

Quantitative examination of anaphase spindle distribution (Goldstein et al., 2017) revealed that synthetic variants also regulated the translocation of Cin8 from the SPBs to the midzone (Fig. 4D), as indicated by reduced intensity at the SPBs and elevated intensity at the midzone compared to the phosphodeficient Cin8-5A. At short and intermediate anaphase spindles, the full Cin8_syn_148f variant exhibited higher distribution to the midzone and lower distribution to the SPBs compared to the partial Cin8_syn_148p counterpart (Fig. 4D 3-4 μm and 4-5 μm red arrows). These results indicate that the synthetic Cdk1 sites regulate the localization of Cin8 on anaphase spindles and that this regulation is more pronounced in full sites compared to partial Cdk1 consensus sequences. Finally, the translocation from the SPBs to the spindle of the full Cin8_syn_276f variant was less pronounced compared to the full variants at the S148 site. Taken together, our data indicate that the two novel sites produce different effects, with S276 mainly affecting detachment from the spindle (Fig. 4A, B, EV1, EV2, and EV3), and the S148 site affecting both Cin8 translocation to the midzone at mid anaphase and detachment from the spindle at late anaphase (Figs. 4, EV1, EV2, and EV3).

To examine whether the newly created Cdk1 sites are accessible for phosphorylation by Clb2/Cdk1, we performed *in vitro* phosphorylation assays using a purified Cin8 motor domain as a substrate, as previously described (Avunie-Masala et al., 2011, Goldstein et al., 2017). Our data indicate that all native variants, S277, S285, and S493, undergo phosphorylation *in vitro*. At the S277 and S285 sites, phosphorylation of the full Cdk1 phosphorylation consensuses is ~20-fold higher than of their partial counterparts (Fig. 5 right), consistent with previous reports (Chang, Begum et al., 2007, Koivomagi, Valk et al., 2011b, Loog & Morgan, 2005). These higher phosphorylation levels do not correlate with the relatively small differences, if any, in Cin8 localization patterns, along and perpendicular to the spindle, between the partial and full phosphorylation consensus variants of these positions (Fig. 2). In contrast, variants with Cdk1 consensus at position S148 exhibited a low degree of phosphorylation compared to wt Cin8 and a less than ~2-fold increase in phosphorylation levels between the partial and the full Cdk1 phosphorylation consensus sites. This result indicates that position S148 is less accessible for Cdk1/Clb2 phosphorylation; however, partial and full consensus sites at this position result in completely different phenotypes. We also included the Cin8_syn_168f variant in this assay, which exhibits S/T independent diffusive Cin8 localization and is also non-viable at 26°C (Fig. 3). This mutant exhibited no phosphorylation *in vitro* (Fig. 5 left), indicating that phenotypes exhibited by this mutant are a result of a deleterious mutation that disrupts Cin8 structure and prevents Cin8 from performing its essential roles. In addition, the Cin8_syn_268f mutant was included in the assay, exhibiting a neutral Cin8-5A-like phenotype, and is a few positions upstream to the native Cdk1 phosphorylation at position S277 in the disordered loop 8. This mutant underwent substantial phosphorylation *in vitro*, ~3-fold higher than wt Cin8 (Fig. 5 right); however, no phosphoregulation of Cin8 localization was evident with this variant (Fig. 3), indicating that extensive phosphorylation at this position caused no phosphoregulation. And finally, we also included Cin8_syn_99 partial and full variants, which were temperature sensitive (Fig. 3). These mutants also exhibited high levels of *in vitro* phosphorylation with ~20-fold increase between partial and full Cdk1 sites (Fig. 5, left), but no detachment from the spindle (Fig. 3). Taken together, these results indicate that the precise position of the phosphorylation site, rather than the degree of phosphorylation, is the most important determinant of Cin8 phosphoregulation.

### Synthetic Cdk1 sites control spindle dynamics

Finally, we examined the control of spindle dynamics by Cin8 phospho-variants (Figs. 6 and EV6). Spindle length was measured as a function of time, and rate was determined for the first (fast) and second (slow) phases of spindle elongation, as previously described (Avunie-Masala et al., 2011, Gerson-Gurwitz et al., 2009, Goldstein et al., 2017, Movshovich et al., 2008, Straight et al., 1998). For simplicity, Fig. 6 exhibits comparison of spindle elongation during the two phases between the full Cin8 variants, while Fig. EV6 presents the full comparison including the partial and control counterparts of these sites. Consistent with our previous report (Goldstein et al., 2017), we found no difference in the rate of elongation between the phosphodeficient Cin8-5A and wt Cin8 during the first phase (Fig. 6 left and EV6). In the slow phase, the rate of elongation of the phosphodeficient Cin8-5A variant was slightly faster than that of wt Cin8; however, this difference was not significant according to post-ANOVA comparison with Tukey procedure (Fig. 6 right). We found that the full variants at the native sites have no effect on spindle elongation during either elongation phase compared to wt Cin8 and Cin8-5A. On the other hand, our data indicate that the synthetic variants at positions S148 and S276 did alter anaphase spindle dynamics. The synthetic variant at position S148 significantly reduced the rate of elongation during the early fast elongation compared to the full native variants Cin8_nat_277 and Cin8_nat_285 (Fig. 6 left), while the full synthetic variant at position S276 significantly reduced the elongation rate during the slower second phase compared to the full native sites Cin8_nat_277 and Cin8_nat_285 (Fig. 6 right), with no effect on the fast phase. This suggests that whereas single native sites bearing full Cdk1 phosphorylation sites have little to no effect on spindle elongation rate, the synthetic sites are affecting Cin8 function during spindle elongation. Interestingly, although the synthetic site S276 is in very high proximity to the native site S277, their regulation of spindle elongation is very different (Fig. 6 right). Examination of rates of elongation of the partial and control variant also demonstrate a complex picture consistent with the notion that the synthetic variants differ from the native sites (Fig. EV6). For example, the variants bearing full Cdk1 phosphorylation consensus at native sites, Cin8_nat_277f and Cin8_nat_285f, exhibit faster elongation rates during the first phase compared to their partial counterparts, with control mutations having similar elevated rates (Fig. EV6 top). In contrast, the rate of the full variant Cin8_syn_148f is not faster than the partial variants and the control variants are significantly faster compared to the full variants (Fig. EV6 top). These results indicate that the synthetic phospho-variants affect spindle elongation in a unique manner, different from that of the native variants, likely affecting spindle localization and/or motor activity by mechanisms different from those of wt Cin8.

## Discussion

In this study we examined the plasticity of phosphoregulation of the Cin8 mitotic kinesin-5 motor by Cdk1 in *S. cerevisiae* cells. We employed an extensive synthetic mutation experiment and characterized the functionality of Cin8 harboring both native and synthetic sites by four *in vivo* assays: viability test, translocation to the midzone in mid anaphase, release from the spindle in late anaphase, and rate of anaphase spindle elongation. We revealed the complexity of the effect of phosphorylation consensus at native Cdk1 sites and found two novel sites that conferred phosphoregulation, which did not, however, fully recapitulate the original phosphoregulation.

### Possible mechanisms of phosphoregulation of Cin8 by native and synthetic Cdk1 sites

Most Cdk1 substrates contain multiple phosphorylation sites (Holt et al., 2009, Moses, Heriche et al., 2007). To initiate this study, we first investigated to what extent single sites were able to confer regulation. We found that each of the three native Cdk1 sites in the catalytic domain of Cin8 were functional alone. The three phosphorylation sites together could give more robust regulation, or perhaps fine-tune the activity of Cin8. Indeed, not all sites are exactly equal and, consistent with our previous work (Goldstein et al., 2017), there is evidence of cooperative regulation of Cin8. A number of kinesin-related proteins were shown to be regulated by phosphorylation at residues outside the catalytic domain (Andrews, Ovechkina et al., 2004, Bishop, Han et al., 2005, Blangy et al., 1995, Cahu et al., 2008, Drummond & Hagan, 1998, Fu, Ward et al., 2009, Giet, Uzbekov et al., 1999, Matsuoka, Ballif et al., 2007, Morfini, Szebenyi et al., 2002, Olsen, Vermeulen et al., 2010, Pigino, Morfini et al., 2009, Rapley, Nicolas et al., 2008, Sawin & Mitchison, 1995, Sharp et al., 1999, Smith, Hegarat et al., 2011, Wojcik et al., 2013, Zhang, Shao et al., 2011, Zhang, Ems-McClung et al., 2008). However, some kinesin motors, Cin8 included, were shown to be regulated by phosphorylation in their catalytic domain. In such cases, phosphorylation affected the motile properties of kinesin and their interaction with MTs (Avunie-Masala et al., 2011, Gerson-Gurwitz, Thiede et al., 2011, Hara & Kimura, 2009, Hizlan, Mishima et al., 2006, Mennella, Tan et al., 2009, Padzik, Deshpande et al., 2016, Shapira & Gheber, 2016), indicating that phosphorylation directly regulates the kinesin catalytic cycle. Thus, it is possible that the synthetic Cdk1 sites in the catalytic domain of Cin8 that confer phosphoregulation, directly affect its catalytic activity.

The native S493 site is located in loop 14 and is conserved among the kinesin-5 homologs (Fig. 7A). This site is located in the vicinity of the Cin8 ATP-binding pocket (Fig. 8), raising the possibility that phosphorylation may affect ATPase activity. The two other native sites, S277 and T285, are located in loop 8 of Cin8, which contains an unusually large insertion not present in the paralogous motor Kip1 and metazoan kinesin-5 motors. In the MT-bound state, loop 8 is located near the MT lattice (Cao, Wang et al., 2014, Goulet & Moores, 2013, Ogawa, Saijo et al., 2017, Wang, Cantos-Fernandes et al., 2017). Loop 8 was shown to regulate the directionality of Cin8 (Gerson-Gurwitz et al., 2011, Shapira & Gheber, 2016) and the non-canonical binding of Cin8 to MTs (Bell, Cha et al., 2017). One of the synthetic phospho-variants, S276, is located in loop 8, one amino-acid apart from the native S277 site (Fig. 7B). The phenotypes of this site resemble the phenotypes of the wt Cin8 and of the native Cin8_nat_277 variant (Figs. 2, 3, 4, and EV5), although they differ in in spindle elongation dynamics (Fig. 6), indicating that the mechanisms of regulation of the S276 and S277 sites are similar, but not identical. Two additional synthetic sites, Cin8_syn_286 and Cin8_syn_300 were introduced into loop 8, and one of these was efficiently phosphorylated *in vitro*, but these sites conferred no regulation (Fig. 3). It is possible that phosphorylation within loop 8, at the S277_nat_ and S276_syn_ positions, changes the conformation of loop 8, thus altering its interaction with the MT lattice and affecting the function of Cin8 and this conformational change requires a precise coordination of the phosphate.

**Figure 7:**
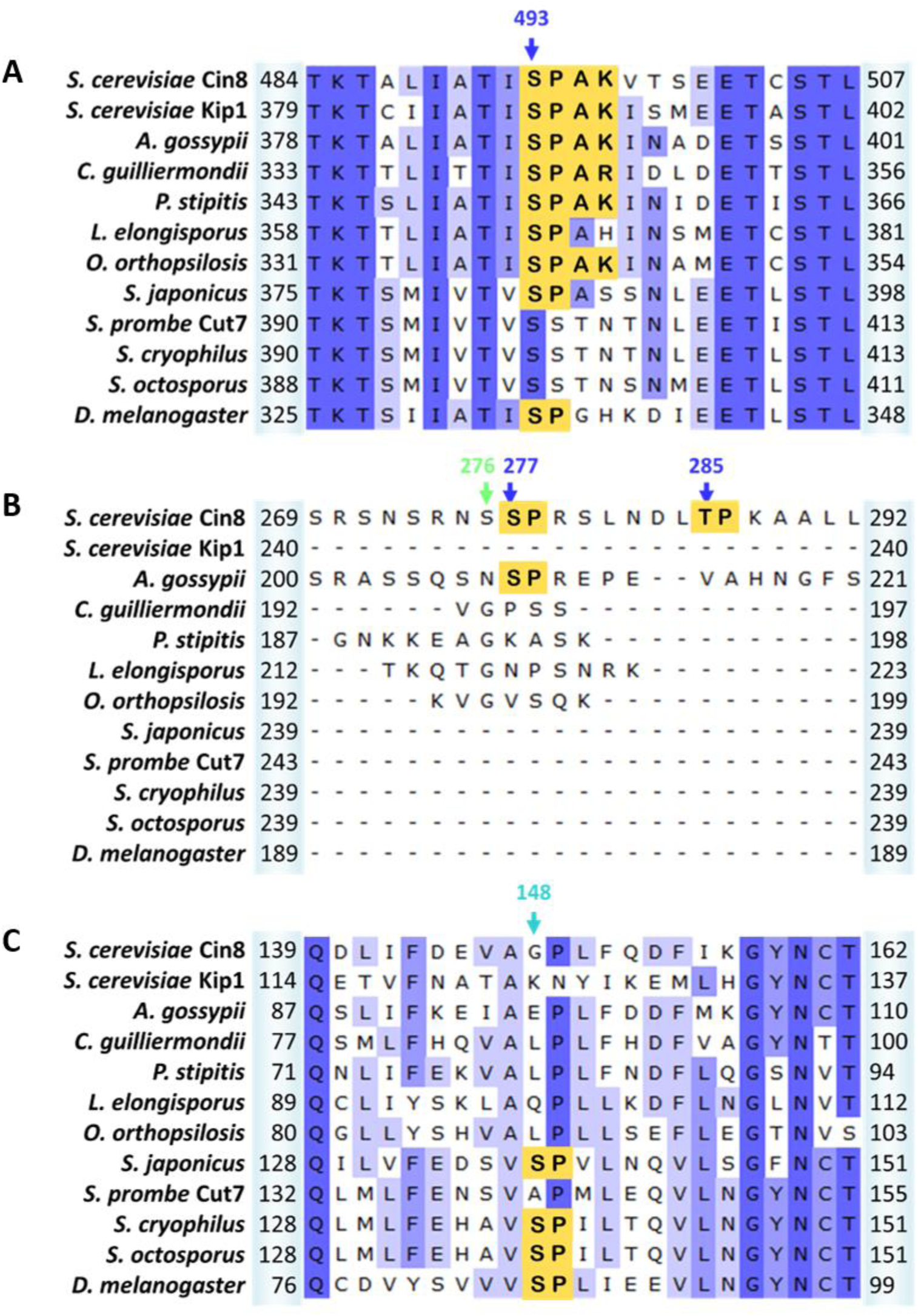
The functional synthetic phosphorylation sites are near to sites that have been sampled in evolution. Multiple sequence alignment of Cin8 with other kinesin-5 proteins using Unipro UGENE alignment tool. Organisms are indicated on the left and listed in the materials and methods section; amino acid positions are indicated on each side of each sequence. A, B, and C demonstrate alignment in the regions near native site 493, native sites in loop 8 and 276, and position 148 in Cin8, respectively. Cdk1 phosphorylation consensuses are highlighted in orange.

**Figure 8:**
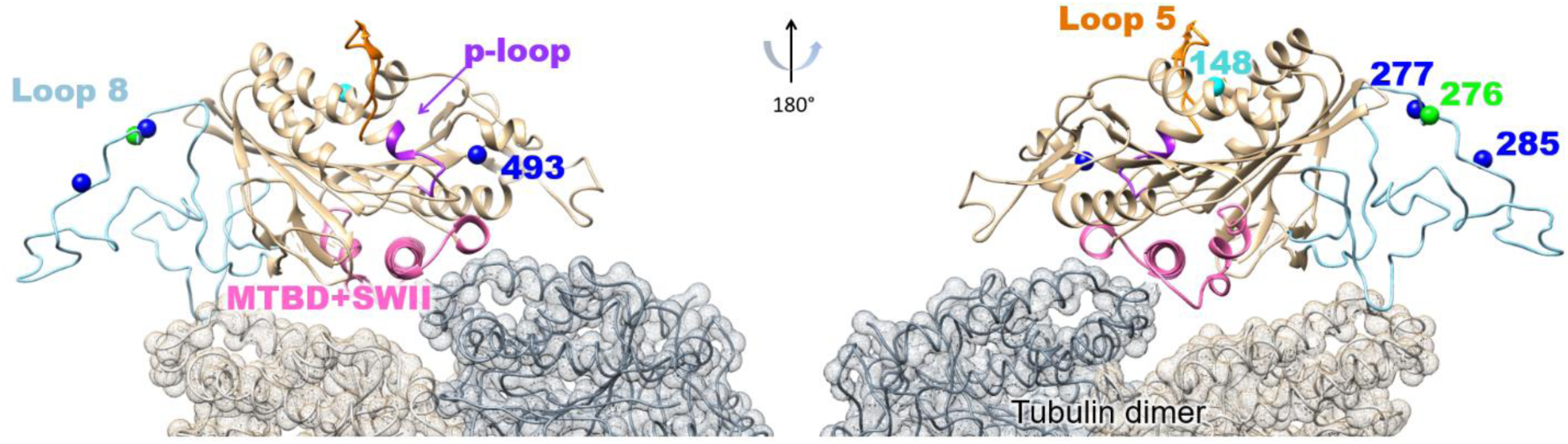
Location of the synthetic phosphorylation sites. Model of Cin8 motor domain in complex with αβ-tubulin predicted by Swiss-model based on pseudo-atomic model of MT-bound *S. pombe* kinesin-5 motor domain in the AMPPNP state (PDB ID: 5M5I) facing in two different orientations. Key-role domains are color-coded; ATPase p-loop in purple (loop 4 with first 3 residues of helix2), MT binding domain and switch II suffixed by “MTDB+SWII” (loop 11, helix 4, loop 12 and helix 5) in pink loop-5 in orange (Kull & Endow, 2002, Sablin, Kull et al., 1996, Subbiah & Harrison, 1989); native Cdk1 phosphorylation sites are represented as spheres: native sites are colored in blue, and synthetic phospho-variants at positions 276 and 148 are colored in light green and cyan, respectively.

Sequence alignment reveals that the second functional synthetic site, Cin8_syn_148, is also found in some kinesin-5 homologs (Fig. 7C), suggesting that this is an important site for kinesin-5 regulation that has been employed elsewhere in evolution. This site is located in α- helix 1 (Subbiah & Harrison, 1989) in the vicinity of loop 5 (Fig. 8) within the kinesin motor domain (Harrington, Naber et al., 2011, Maliga, Xing et al., 2006, Waitzman, Larson et al., 2011). Although the function of loop 5 is unknown, its proximity to the conserved nucleotide-binding P-loop element and the fact that several kinesin-5 inhibitors bind to this loop (Brier, Lemaire et al., 2004, Lad, Luo et al., 2008) indicate that loop 5 is important for kinesin-5 function. Thus, phosphorylation of the S148 site may affect the function of kinesin-5 motors by affecting the conformation of loop 5.

### Correlation between in vitro phosphorylation and *in vivo* phosphoregulation

Results presented here indicate that the two synthetic variants at positions S148 and S276 affected spindle elongation rates in a phosphorylation-dependent manner (Fig. EV6). Changing from a partial to full consensus Cdk1 sequence decreased the elongation rate of the S148 variant, and introducing a non-phosphorylatable control mutation fully rescued this effect (Fig. EV6). This result suggests that synthetic phosphoregulation sites can modulate the dynamics of the spindle to various degrees *in vivo*. Thus, a single synthetic phosphoregulatory site can tune intracellular functions.

Previous results have indicated that phosphorylation of the three native sites in the catalytic domain of Cin8 is required for its detachment from the spindle (Avunie-Masala et al., 2011). However, results presented here indicate that there is only partial correlation between *in vitro* phosphorylation and *in vivo* spindle detachment of the different phospho-variants. For example, the synthetic variants at positions S99 and S268 exhibited a high level of phosphorylation *in vitro* that was considerably higher than phosphorylation of wt Cin8 or single native variants (Fig. 5, right). However, in spite of this high level of phosphorylation the synthetic variants don’t detach from the spindle at late anaphase, but rather exhibit a Cin8-5A-like phenotype (Fig. 3). In contrast, the full variant at position S148 exhibits a very low level of phosphorylation *in vitro* (Fig. 5 left), and yet exhibits high levels of detachment from the spindle (Figs. 3, 4B, EV1, EV2, and EV3). These results clearly indicate that although phosphorylation at a single amino-acid can induce detachment from the spindle, as with the native variants, a high level of phosphorylation in itself is insufficient to induce spindle detachment. Thus, the precise location of phosphorylation plays a critical role in phosphoregulation of Cin8.

### Cell cycle networks are flexible, Cin8 regulation is constrained

To date there has been very little experimental data systematically testing the flexibility of phosphoregulation. Cell cycle networks have been extensively rewired over long evolutionary time-scales. For example, while the general cell-cycle network topologies are conserved from yeast to humans, many of the molecular components have been replaced (Cross, Buchler et al., 2011). Thus, it is clear that there is considerable flexibility in cell cycle regulation. This general feature is likely related to the fact that >90% of Cdk1-dependent phosphorylation events occur in disordered regions of proteins (Holt et al., 2009). Nevertheless, there is clearly constraint on phosphorylation sites (Nguyen Ba & Moses, 2010). As a number of kinesin-related proteins have been shown to be regulated by phosphorylation at different regions of the molecule we chose to investigate the flexibility of phosphoregulation using a mitotic kinesin-5 Cin8 as a model.

We found that two out of 29 Cin8_syn_ mutations resulted in phosphorylation-dependent detachments. Of these two, one (position S276) was viable at all temperatures. Furthermore, although the phenotypes of the Cin8_syn_276 variant were similar to those of the native Cin8_nat_277 sites, the two alleles differed in their distribution along the spindle (Fig. EV5, left), timing of their detachment from the spindle (Fig. EV5, right), and in rate of anaphase spindle elongation (Fig. 6). Thus, although some of the aspects of phosphoregulation at the synthetic sites were similar to those of the native sites, none of the new synthetic sites precisely recapitulated the phenotypes of the native sites.

In summary, the data presented here indicate that phosphoregulation of the kineisn-5 Cin8 by Cdk1 is site-specific and constrained. A synthetic Cdk1 site only one amino-acid distant from a native site within a disordered loop could not fully recapitulate the native regulation. Kinesin-5 motors function by binding and moving along MTs via a precise catalytic cycle, with directionality, motor activity, and spindle binding all subject to phosphoregulation. Thus, regulation must be highly constrained to maintain optimal control of mitotic kinesin motors such that they can efficiently perform their role in mitotic spindle dynamics.

## Materials and Methods

### Generation of synthetic Cdk1 phospho-variants and their categorization

The synthetic Cdk1 sites of Cin8 were generated on the basis of the phosphodeficient variant of Cin8 in which serines or threonines in all five native Cdk1 sites (S277, T285, S493, S763, and S1010) were mutated to alanine. Sites for novel synthetic Cdk1 phospho-variants were chosen in a systematic manner, with each original proline in the sequence of Cin8 being targeted as a potential Cdk1 phosphorylation site. Out of 20 prolines in the sequence of full length Cin8, five are occupied by the native sites and the remaining fifteen prolines were tested for potential Cdk1 phosphoregulation sites. In these cases, to create a Cdk1 site, a serine was added replacing an amino-acid at position -1 to the proline, resulting in a partial (**[S/T]**-P) or full (**[S/T]**-P-x-[K/R]) Cdk1 phosphorylation site (Fig. 1B class 1). In one case, at position 465, a threonine was added at position -1 to an existing proline. In the case where a partial Cdk1 phosphorylation site is created by this mutagenesis, a sequential mutation introduced a lysine at position +3 to the serine to result in a full Cdk1 phosphorylation consensus ([S/T]-P-x-[K/R]) (Fig. 1B class 1). In addition, if an [S/T]-x-x-[K/R] sequence is present in a non-coiled coil region of Cin8, a proline is introduced at position +1 to the serine/threonine, resulting in a full Cdk1 phosphorylation consensus [S/T]-**P**-x-[K/R] (Fig. 1B class 2). The addition of serine at position -1 to the proline and the addition of lysine at position +3 to the serine were preferred over threonine and arginine, respectively, since it results in a stronger phosphorylation consensus (Chang et al., 2007, Koivomagi et al., 2011b, Loog & Morgan, 2005). By this strategy, three of the five native sites, at positions 277, 285, and 493, were converted to [S-P] sites although position 285 originally contained a threonine followed by a proline [T-P]. The two native sites outside of the motor domain, at positions 736 and 1010, were not sampled since they were previously shown to have no phosphoregulation properties (Avunie-Masala et al., 2011). Two sites of class 2, at positions 712 and 271, were not sampled due to technical reasons. Overall, 29 new Cdk1 sites were generated. The novel sites were examined by live cell imaging for possible anaphase spindle-detachment phenotypes (Avunie-Masala et al., 2011, Goldstein et al., 2017). Mutants that were viable at 26°C and exhibited no detachment of Cin8 from the spindle were categorized as Cin8-5A-like mutants, to indicate they are not phosphoregulated similarly to wt Cin8 (colored orange in Fig. 3). Typically, Cin8-5A-like mutants were viable at all temperatures with five exceptions: (a) Cin8_syn_39, which exhibits slight temperature sensitivity at 35°C; (b) Cin8_syn_99 partial and full variants, which exhibit temperature sensitivity, which is partially rescued by the control counterparts; (c) Cin8_syn_103 and Cin8_syn_99 are exhibiting reduced viability at 33°C and 35°C; (d) Cin8_syn_700, which exhibits reduced viability of the full variant but not of the partial variant; and (e) Cin8_syn_881, which exhibits temperature sensitivity at 33°C and 35°C. Since these variants did not detach from the spindles, they were classified as Cin8-5A-like variants and not examined further by quantitative analyses.

Mutants that exhibited detachment from the spindle were targeted as possible Cdk1 phosphoregulation sites and additional control mutants were generated at these positions to determine if the observed phenotypes were phosphorylation-dependent. These control mutants contained a phosphodeficient alanine replacing the serine or threonine within the Cdk1 consensus sequence. In addition, the variants were examined by yeast viability test to assess their ability to function in cells as a sole source of kinesin-5 activity (see below). Eight of the variants that exhibited detachment from the spindles at anaphase were not viable at 26^°^C as full Cdk1 sites (colored in black in Fig. 3). One site in this category, at position 134, was non-viable as a partial Cdk1 phosphorylation consensus of its control variant (Cin8_syn_134pc) partially rescues the phenotype. Since the full variant at this position (Cin8_syn_134f) is non-viable, this site was not examined further by quantitative analysis. Another exception is at position 194, in which the partial variant Cin8_syn_194p was viable at 26^°^C and 33^°^C but temperature sensitive at 35^°^C, with no detachments from the spindle at 26^°^C. However, its full counterpart was non-viable at 26°C, which was not rescued by the control mutant and therefore was categorized as a non-viable variant.

Variants that exhibited phosphorylation-dependent spindle localization phenotypes, at positions S148 and S276, were further investigated by quantitative analysis of their distribution along the spindle and detachments.

#### Yeast viability test

Viability test of cells expressing the Cin8_nat_ and Cin8_syn_ variants as a sole source of kinesin-5 function was performed as previously described (Avunie-Masala et al., 2011, Duselder et al., 2015, Goldstein et al., 2017). Yeast strains used for this assay were deleted for their chromosomal copies of *CIN8* and *KIP1* and contained an endogenic recessive cycloheximide resistance gene (*cin8Δkip1Δcyh^r^*). These cells were supplemented with a plasmid (pMA1208) encoding for wt Cin8 and a wt cycloheximide sensitivity gene. After transformation with a plasmid encoding the Cin8 variant of interest, the initial pMA1208 plasmid was shuffled out by growth on YPD medium containing 7.5 μg/mL cycloheximide for 3 to 4 days, at 26^°^C and elevated temperatures, in a serial dilution (1:1, 1:10, 1:10^2^, 1:10^3^, 1:10^4^) starting with 0.2 O.D_λ=600nm_ (approximately 2.55×10^6^ cells/ml).

#### *In vitro* phosphorylation assay

*In vitro* phosphorylation assays were performed as previously described (Avunie-Masala et al., 2011, Goldstein et al., 2017). In brief, bacterially expressed Cin8 (590)-TEV-EGFP-6His variants were purified using standard nickel affinity chromatography, eluted with 300mM imidazole that was subsequently removed using Zeba Spin Desalting Columns 40K (Thermo Scientific). For a phosphorylation assay, equal concentrations of Cin8 variants were mixed with TAP-purified Clb2-Cdk1-Cks1 complex in kinase assay mixture [50mM HEPES, pH 7.4, 150mM NaCl, 5mM MgCl_2_, 8% glycerol, 0.2 mg/ml BSA, 500nM Cks1, and 500μM ATP (with added γ-^32^P-ATP (PerkinElmer)]. Reactions were stopped after 10 and 20 min with SDS-PAGE sample buffer and proteins were separated by SDS-PAGE. Gels were stained with Coomassie Brilliant Blue (CBB) R-250 (Sigma) and incorporation of ^32^P into the proteins was visualized by autoradiography.

#### Live cell imaging

The *S. cerevisiae* strains and plasmids used in this study are described in SI Tables S1, S2, and S3. Live cell imaging was performed as previously described (Avunie-Masala et al., 2011, Gerson-Gurwitz et al., 2009, Movshovich et al., 2008). Images were acquired using a Zeiss Axiovert 200M-based microscope setup equipped with a cooled CCD Andor Neo sCMOS camera. Images of Z stacks of eleven planes were obtained in three channels with 0.5 μm separation. Time-lapse images were obtained using a Zeiss Axiovert 200M-based Nipkow spinning-disc confocal microscope (UltraView ESR, Perkin Elmer, UK) with an EMCCD camera. Z stacks of 32-36 slices with 0.2 μm separation were acquired at one-minute intervals for 70 minutes. Data analysis was performed using MetaMorph (MDS Analytical Technologies) and open source ImageJ software. **Spindle elongation rates** were determined as previously described (Avunie-Masala et al., 2011).

#### Fluorescence intensity distribution of Cin8 along the spindle

Line scan analysis along the spindle was employed to quantify the distribution of 3GFP-tagged Cin8 variants, as previously described (Fridman et al., 2013, Goldstein et al., 2017). The fluorescence intensity profile was determined along a line tracing the spindle from mother to bud. The background signal was calculated by averaging the intensity outside the nucleus and that value was subtracted from the fluorescence intensity measured at each point. Finally, intensity was interpolated and divided into 100 segments of equal length using Origin software (OriginLab). Normalization of the Cin8-3GFP fluorescent signals was performed by dividing the intensity at each point by the total Cin8-3GFP fluorescence intensity at each spindle (Figs. 2C and 4D). The average intensity was calculated for each length-point for 10-20 cells.

#### Cin8 detachment from the spindle determined from fluorescence intensity perpendicular to the spindle

Line scan analysis perpendicular to the spindle was performed to quantify the detachment of Cin8 variants from the spindle during anaphase as previously described (Goldstein et al., 2017). First, the Cin8-3GFP fluorescence profile was measured along a line of 40 pixels (5.12 μm) drawn perpendicular to the spindle. The center of this line was set 2 pixels towards the midzone from a point of highest intensity, usually near the SPB (Figs. 2D). Eleven outer pixels on each side were assigned as background (Fig. 2D, dashed dark grey). The intensity of 5 pixels in the center of the line was assigned as the intensity near the SPB or the spindle and was not considered in calculating Cin8 detachment (Fig. 2D, dashed light grey). The intensities of pixels 12-18 and 23-29 were considered as the intensity resulting from Cin8-3GFP being localized in the nucleus due to its detachment from the spindle (Fig. 2D, green). Following subtraction of the background, the intensity along the line perpendicular to the spindle was normalized to the highest intensity value, usually observed near the SPB. Then, the intensity of Cin8-3GFP detached from the spindle (pixels 12-18 and 23-29) was averaged for each cell. Finally, the average Cin8-GFP intensity perpendicular to the spindles of 6-7 μm, was calculated in all cells expressing the same variant (10-20 cells – one exception is Cin8_syn_148pc in which only four cells were analyzed). Cin8 detachment from the spindle was analyzed as a function of spindle length (Figs. 2E and 4B) and as a function of time, based on time-lapse images as in Fig. EV1 (Figs. EV3 and EV5 right).

**Statistical analysis** was done using post-ANOVA all-pairwise comparison performed with Tukey procedure using OriginLab software.

#### Multiple sequence alignments (MSAs)

MSAs presented in Fig. 8 were conducted according to a phylogenetic tree as in (Wojcik et al., 2013). Strains chosen (as presented from top to bottom in Fig. 8): Cin8 and Kip1 of *Saccharomyces cerevisiae, Ashbya gossypii (Eremothecium gossypii), Candida guilliermondii (Meyerozyma guilliermondii), Scheffersomyces stipites, Lodderomyces elongisporus, Candida orthopsilosis, Schizosaccharomyces japonicus, Schizosaccharomyces pombe, Schizosaccharomyces cryophilus, Schizosaccharomyces octosporus*, and *Drosophila melanogaster*. The MSA was calculated by MUSCLE algorithm via UGENE program. Color coded by percentage identity with a 30% threshold.

#### Stability assay at room and elevated temperatures

Yeast strains were grown in 26°C and 33.5°C for five hours before crude protein extraction with 0.1M NaOH. Cell lysate was diluted 1:1 with SDS-PAGE sample-buffer. The samples were separated by SDS-PAGE and blotted by Western blot to a PDVF membrane and developed with α-GFP HRP conjugated anti-body (Santa Cruz GFP Antibody (B-2): sc-9996).

## Acknowledgements

We thank Kumar-Singh Sudhir, Pandey Himanshu, Siegler Nurit, and Popov Mary from Ben-Gurion University in the Negev, Beer-Sheva, Israel, for critical reading of this manuscript. This work was supported in part by the Israel Science Foundation (ISF) (grant 165/13) awarded to L.G.; The United States - Israel Binational Science Foundation grant (BSF-2015851), awarded to L.G.; the William Bowes Foundation and Vilcek Foundation grant awarded to L.J.H.; The ERC Consolidator Grant Nr 649124, Phosphoprocessors, awarded to M.L.; and Estonian Science Agency grant Nr. IUT2-21, awarded to M.L.

## Author contributions

A.G. planned and performed experiments, analyzed data, designed figures, wrote and revised the manuscript. D.G. performed experiments. E.V. performed experiments. M.L. supervised experiments and revised the manuscript. L.J.H. conceived the study, planned experiments, wrote and revised the manuscript. L.G. conceived the study, planned experiments, supervised experiments, wrote and revised the manuscript.

## References

Andrews PD, Ovechkina Y, Morrice N, Wagenbach M, Duncan K, Wordeman L, Swedlow JR (2004) Aurora B regulates MCAK at the mitotic centromere. Dev Cell 6: 253–268

Avunie-Masala R, Movshovich N, Nissenkorn Y, Gerson-Gurwitz A, Fridman V, Koivomagi M, Loog M, Hoyt MA, Zaritsky A, Gheber L (2011) Phospho-regulation of kinesin-5 during anaphase spindle elongation. J Cell Sci 124: 873–878

Bell KM, Cha HK, Sindelar CV, Cochran JC (2017) The yeast kinesin-5 Cin8 interacts with the microtubule in a noncanonical manner. J Biol Chem 12: 797662

Bishop JD, Han Z, Schumacher JM (2005) The Caenorhabditis elegans Aurora B kinase AIR-2 phosphorylates and is required for the localization of a BimC kinesin to meiotic and mitotic spindles. Mol Biol Cell 16: 742–56

Blangy A, Arnaud L, Nigg EA (1997) Phosphorylation by p34cdc2 protein kinase regulates binding of the kinesin-related motor HsEg5 to the dynactin subunit p150. J Biol Chem 272: 19418–19424

Blangy A, Lane HA, d’Herin P, Harper M, Kress M, Nigg EA (1995) Phosphorylation by p34cdc2 regulates spindle association of human Eg5, a kinesin-related motor essential for bipolar spindle formation in vivo. Cell 83: 1159–69

Brier S, Lemaire D, Debonis S, Forest E, Kozielski F (2004) Identification of the protein binding region of S-trityl-L-cysteine, a new potent inhibitor of the mitotic kinesin Eg5. Biochemistry 43: 13072–82

Cahu J, Olichon A, Hentrich C, Schek H, Drinjakovic J, Zhang C, Doherty-Kirby A, Lajoie G, Surrey T (2008) Phosphorylation by Cdk1 increases the binding of Eg5 to microtubules in vitro and in Xenopus egg extract spindles. PLoS ONE 3: e3936

Cao L, Wang W, Jiang Q, Wang C, Knossow M, Gigant B (2014) The structure of apo-kinesin bound to tubulin links the nucleotide cycle to movement. Nat Commun 5: 5364

Chang EJ, Begum R, Chait BT, Gaasterland T (2007) Prediction of cyclin-dependent kinase phosphorylation substrates. PLoS One 2: e656

Chee MK, Haase SB (2010) B-cyclin/CDKs regulate mitotic spindle assembly by phosphorylating kinesins-5 in budding yeast. PLoS Genet 6: e1000935

Crasta K, Huang P Fau - Morgan G, Morgan G Fau - Winey M, Winey M Fau - Surana U, Surana U (2006) Cdk1 regulates centrosome separation by restraining proteolysis of microtubule-associated proteins.

Cross FR, Buchler NE, Skotheim JM (2011) Evolution of networks and sequences in eukaryotic cell cycle control. Philos Trans R Soc Lond B Biol Sci 366: 3532–44

D’Amours D, Amon A (2004) At the interface between signaling and executing anaphase-- Cdc14 and the FEAR network. Genes Dev 18: 2581–95

Drummond DR, Hagan IM (1998) Mutations in the bimC box of Cut7 indicate divergence of regulation within the bimC family of kinesin related proteins. J Cell Sci 111 (Pt 7): 853–65

Duselder A, Fridman V, Thiede C, Wiesbaum A, Goldstein A, Klopfenstein DR, Zaitseva O, Janson ME, Gheber L, Schmidt CF (2015) Deletion of the Tail Domain of the Kinesin-5 Cin8 affects its directionality. J Biol Chem 19: 620799

Ferenz NP, Gable A, Wadsworth P (2010) Mitotic functions of kinesin-5. Semin Cell Dev Biol 21: 255–9

Fridman V, Gerson-Gurwitz A, Movshovich N, Kupiec M, Gheber L (2009) Midzone organization restricts interpolar microtubule plus-end dynamics during spindle elongation. EMBO Rep 10: 387–93

Fridman V, Gerson-Gurwitz A, Shapira O, Movshovich N, Lakamper S, Schmidt CF, Gheber L (2013) Kinesin-5 Kip1 is a bi-directional motor that stabilizes microtubules and tracks their plus-ends in vivo. J Cell Sci 126: 4147–59

Fu C, Ward JJ, Loiodice I, Velve-Casquillas G, Nedelec FJ, Tran PT (2009) Phospho-regulated interaction between kinesin-6 Klp9p and microtubule bundler Ase1p promotes spindle elongation. Dev Cell 17: 257–67

Gardner MK, Bouck DC, Paliulis LV, Meehl JB, O’Toole ET, Haase J, Soubry A, Joglekar AP, Winey M, Salmon ED, Bloom K, Odde DJ (2008) Chromosome congression by Kinesin-5 motor-mediated disassembly of longer kinetochore microtubules. Cell 135: 894–906

Gerson-Gurwitz A, Movshovich N, Avunie R, Fridman V, Moyal K, Katz B, Hoyt MA, Gheber L (2009) Mid-anaphase arrest in S. cerevisiae cells eliminated for the function of Cin8 and dynein. Cell Mol Life Sci 66: 301–13

Gerson-Gurwitz A, Thiede C, Movshovich N, Fridman V, Podolskaya M, Danieli T, Lakamper S, Klopfenstein DR, Schmidt CF, Gheber L (2011) Directionality of individual kinesin-5 Cin8 motors is modulated by loop 8, ionic strength and microtubule geometry. Embo J 30: 4942–54

Gheber L, Kuo SC, Hoyt MA (1999) Motile properties of the kinesin-related Cin8p spindle motor extracted from Saccharomyces cerevisiae cells. J Biol Chem 274: 9564–72

Gibbs KL, Greensmith L, Schiavo G (2015) Regulation of Axonal Transport by Protein Kinases. Trends Biochem Sci 40: 597–610

Giet R, Uzbekov R, Cubizolles F, Le Guellec K, Prigent C (1999) The Xenopus laevis aurora-related protein kinase pEg2 associates with and phosphorylates the kinesin-related protein XlEg5. J Biol Chem 274: 15005–15013

Goldstein A, Siegler N, Goldman D, Judah H, Valk E, Koivomagi M, Loog M, Gheber L (2017) Three Cdk1 sites in the kinesin-5 Cin8 catalytic domain coordinate motor localization and activity during anaphase. Cell Mol Life Sci 74(18):3395–3412.

Goulet A, Moores C (2013) New insights into the mechanism of force generation by kinesin-5 molecular motors. Int Rev Cell Mol Biol 304: 419–66

Hara Y, Kimura A (2009) Cell-size-dependent spindle elongation in the Caenorhabditis elegans early embryo. Curr Biol 19: 1549–54

Harrington TD, Naber N, Larson AG, Cooke R, Rice SE, Pate E (2011) Analysis of the interaction of the Eg5 Loop5 with the nucleotide site. J Theor Biol 289: 107–15

Hildebrandt ER, Hoyt MA (2000) Mitotic motors in Saccharomyces cerevisiae. Biochim Biophys Acta 1496: 99–116.

Hizlan D, Mishima M, Tittmann P, Gross H, Glotzer M, Hoenger A (2006) Structural analysis of the ZEN-4/CeMKLP1 motor domain and its interaction with microtubules. J Struct Biol 153: 73–84

Holt LJ, Tuch BB, Villen J, Johnson AD, Gygi SP, Morgan DO (2009) Global analysis of Cdk1 substrate phosphorylation sites provides insights into evolution. Science 325: 1682–6

Hoyt MA, He L, Loo KK, Saunders WS (1992) Two Saccharomyces cerevisiae kinesin-related gene products required for mitotic spindle assembly. J Cell Biol 118: 109–20

Ibarlucea-Benitez I, Ferro LS, Drubin DG, Barnes G (2018) Kinesins relocalize the chromosomal passenger complex to the midzone for spindle disassembly. J Cell Biol

Kashina AS, Rogers GC, Scholey JM (1997) The bimC family of kinesins: essential bipolar mitotic motors driving centrosome separation. Biochim Biophys Acta 1357: 257–71

Koivomagi M, Ord M, Iofik A, Valk E, Venta R, Faustova I, Kivi R, Balog ER, Rubin SM, Loog M (2013) Multisite phosphorylation networks as signal processors for Cdk1. Nat Struct Mol Biol 20: 1415–24

Koivomagi M, Valk E, Venta R, Iofik A, Lepiku M, Balog ER, Rubin SM, Morgan DO, Loog M (2011a) Cascades of multisite phosphorylation control Sic1 destruction at the onset of S phase. Nature 480(7375):128–131.

Koivomagi M, Valk E, Venta R, Iofik A, Lepiku M, Morgan DO, Loog M (2011b) Dynamics of Cdk1 substrate specificity during the cell cycle. Mol Cell 42: 610–23

Kull FJ, Endow SA (2002) Kinesin: switch I & II and the motor mechanism. J Cell Sci 115: 15–23

Lad L, Luo L, Carson JD, Wood KW, Hartman JJ, Copeland RA, Sakowicz R (2008) Mechanism of inhibition of human KSP by ispinesib. Biochemistry 47: 3576–85

Loog M, Morgan DO (2005) Cyclin specificity in the phosphorylation of cyclin-dependent kinase substrates. Nature 434: 104–8

Maliga Z, Xing J, Cheung H, Juszczak LJ, Friedman JM, Rosenfeld SS (2006) A pathway of structural changes produced by monastrol binding to Eg5. J Biol Chem 281: 7977–82

Matsuoka S, Ballif BA, Smogorzewska A, McDonald ER, 3rd, Hurov KE, Luo J, Bakalarski CE, Zhao Z, Solimini N, Lerenthal Y, Shiloh Y, Gygi SP, Elledge SJ (2007) ATM and ATR substrate analysis reveals extensive protein networks responsive to DNA damage. Science 316: 1160–6

Mendenhall MD, Jones CA, Reed SI (1987) Dual regulation of the yeast CDC28-p40 protein kinase complex: cell cycle, pheromone, and nutrient limitation effects. Cell 50: 927–35

Mennella V, Tan DY, Buster DW, Asenjo AB, Rath U, Ma A, Sosa HJ, Sharp DJ (2009) Motor domain phosphorylation and regulation of the Drosophila kinesin 13, KLP10A. J Cell Biol 186: 481–90

Morfini G, Schmidt N, Weissmann C, Pigino G, Kins S (2016) Conventional kinesin: Biochemical heterogeneity and functional implications in health and disease. Brain Res Bull 126(Pt 3):347–353.

Morfini G, Szebenyi G, Elluru R, Ratner N, Brady ST (2002) Glycogen synthase kinase 3 phosphorylates kinesin light chains and negatively regulates kinesin-based motility. EMBO J 21: 281–93

Morgan DO, Roberts JM (2002) Oscillation sensation. Nature 418: 495–6

Moses AM, Heriche JK, Durbin R (2007) Clustering of phosphorylation site recognition motifs can be exploited to predict the targets of cyclin-dependent kinase. Genome Biol 8: R23

Movshovich N, Fridman V, Gerson-Gurwitz A, Shumacher I, Gertsberg I, Fich A, Hoyt MA, Katz B, Gheber L (2008) Slk19-dependent mid-anaphase pause in kinesin-5-mutated cells. J Cell Sci 121: 2529–39

Nasmyth K (1996) At the heart of the budding yeast cell cycle. Trends Genet 12: 405–12

Nguyen Ba AN, Moses AM (2010) Evolution of characterized phosphorylation sites in budding yeast. Mol Biol Evol 27: 2027–37

Ogawa T, Saijo S, Shimizu N, Jiang X, Hirokawa N (2017) Mechanism of Catalytic Microtubule Depolymerization via KIF2-Tubulin Transitional Conformation. Cell Rep 20: 2626–2638

Olsen JV, Vermeulen M, Santamaria A, Kumar C, Miller ML, Jensen LJ, Gnad F, Cox J, Jensen TS, Nigg EA, Brunak S, Mann M (2010) Quantitative phosphoproteomics reveals widespread full phosphorylation site occupancy during mitosis. Sci Signal 3: ra3

Padzik A, Deshpande P, Hollos P, Franker M, Rannikko EH, Cai D, Prus P, Magard M, Westerlund N, Verhey KJ, James P, Hoogenraad CC, Coffey ET (2016) KIF5C S176 Phosphorylation Regulates Microtubule Binding and Transport Efficiency in Mammalian Neurons. Front Cell Neurosci 10: 57

Pigino G, Morfini G, Atagi Y, Deshpande A, Yu C, Jungbauer L, LaDu M, Busciglio J, Brady S (2009) Disruption of fast axonal transport is a pathogenic mechanism for intraneuronal amyloid beta. Proc Natl Acad Sci U S A 106: 5907–12

Queralt E, Lehane C, Novak B, Uhlmann F (2006) Downregulation of PP2A(Cdc55) phosphatase by separase initiates mitotic exit in budding yeast. Cell 125: 719–32

Rapley J, Nicolas M, Groen A, Regue L, Bertran MT, Caelles C, Avruch J, Roig J (2008) The NIMA-family kinase Nek6 phosphorylates the kinesin Eg5 at a novel site necessary for mitotic spindle formation. J Cell Sci 121: 3912–21

Ritter A, Kreis NN, Louwen F, Wordeman L, Yuan J (2015) Molecular insight into the regulation and function of MCAK. Crit Rev Biochem Mol Biol 51: 228–45

Roof DM, Meluh PB, Rose MD (1992) Kinesin-related proteins required for assembly of the mitotic spindle. J Cell Biol 118: 95–108

Sablin EP, Kull FJ, Cooke R, Vale RD, Fletterick RJ (1996) Crystal structure of the motor domain of the kinesin-related motor ncd. Nature 380: 555–559

Saunders WS, Hoyt MA (1992) Kinesin-related proteins required for structural integrity of the mitotic spindle. Cell 70: 451–8

Saunders WS, Koshland D, Eshel D, Gibbons IR, Hoyt MA (1995) Saccharomyces cerevisiae kinesin- and dynein-related proteins required for anaphase chromosome segregation. J Cell Biol 128: 617–24

Sawin KE, Mitchison TJ (1995) Mutations in the kinesin-like protein Eg5 disrupting localization to the mitotic spindle. Proc Natl Acad Sci U S A 92: 4289–93

Schwob E, Bohm T, Mendenhall MD, Nasmyth K (1994) The B-type cyclin kinase inhibitor p40SIC1 controls the G1 to S transition in S. cerevisiae. Cell 79: 233–44

Shapira O, Gheber L (2016) Motile properties of the bi-directional kinesin-5 Cin8 are affected by phosphorylation in its motor domain. Sci Rep 6: 25597

Shapira O, Goldstein A, Al-Bassam J, Gheber L (2016) Regulation and Possible Physiological Role of BI-Directional Motility of the Kinesin-5 Cin8. Biophysical Journal 110: 460a

Sharp DJ, McDonald KL, Brown HM, Matthies HJ, Walczak C, Vale RD, Mitchison TJ, Scholey JM (1999) The bipolar kinesin, KLP61F, cross-links microtubules within interpolar microtubule bundles of Drosophila embryonic mitotic spindles. J Cell Biol 144: 125–38

Singh SK, Pandey H, Al-Bassam J, Gheber L (2018) Bidirectional motility of kinesin-5 motor proteins: structural determinants, cumulative functions and physiological roles. Cell Mol Life Sci 75(10):1757–1771.

Smith E, Hegarat N, Vesely C, Roseboom I, Larch C, Streicher H, Straatman K, Flynn H, Skehel M, Hirota T, Kuriyama R, Hochegger H (2011) Differential control of Eg5-dependent centrosome separation by Plk1 and Cdk1. EMBO J 30: 2233–45

Stegmeier F, Amon A (2004) Closing mitosis: the functions of the Cdc14 phosphatase and its regulation. Annu Rev Genet 38: 203–32

Straight AF, Sedat JW, Murray AW (1998) Time-lapse microscopy reveals unique roles for kinesins during anaphase in budding yeast. J Cell Biol 143: 687–94

Subbiah S, Harrison SC (1989) A simulated annealing approach to the search problem of protein crystallography. Acta Crystallogr A 45 ( Pt 5): 337–42

Tytell JD, Sorger PK (2006) Analysis of kinesin motor function at budding yeast kinetochores.JCB 172: 861–874

Valentine MT, Fordyce PM, Block SM (2006) Eg5 steps it up! Cell Div 1: 31

Valentine MT, Gilbert SP (2007) To step or not to step? How biochemistry and mechanics influence processivity in Kinesin and Eg5. Curr Opin Cell Biol 19: 75–81

Waitzman JS, Larson AG, Cochran JC, Naber N, Cooke R, Jon Kull F, Pate E, Rice SE (2011) The loop 5 element structurally and kinetically coordinates dimers of the human kinesin-5, Eg5. Biophys J 101: 2760–9

Waitzman JS, Rice SE (2014) Mechanism and regulation of kinesin-5, an essential motor for the mitotic spindle. Biol Cell 106: 1–12

Wang W, Cantos-Fernandes S, Lv Y, Kuerban H, Ahmad S, Wang C, Gigant B (2017) Insight into microtubule disassembly by kinesin-13s from the structure of Kif2C bound to tubulin. Nat Commun 8: 70

Wargacki MM, Tay JC, Muller EG, Asbury CL, Davis TN (2010) Kip3, the yeast kinesin-8, is required for clustering of kinetochores at metaphase. Cell Cycle 9: 2581–8

Wojcik EJ, Buckley RS, Richard J, Liu L, Huckaba TM, Kim S (2013) Kinesin-5: Cross-bridging mechanism to targeted clinical therapy. Gene 531: 133–49

Zhang L, Shao H, Huang Y, Yan F, Chu Y, Hou H, Zhu M, Fu C, Aikhionbare F, Fang G, Ding X, Yao X (2011) PLK1 phosphorylates mitotic centromere-associated kinesin and promotes its depolymerase activity. J Biol Chem 286: 3033–46

Zhang X, Ems-McClung SC, Walczak CE (2008) Aurora A phosphorylates MCAK to control ran-dependent spindle bipolarity. Mol Biol Cell 19: 2752–65

